# A Patient-Specific Morphoelastic Growth Model of Aortic Dissection Evolution

**DOI:** 10.1101/2024.05.28.596335

**Authors:** Kameel Khabaz, Junsung Kim, Ross Milner, Nhung Nguyen, Luka Pocivavsek

**Affiliations:** David Geffen School of Medicine, University of California, Los Angeles, 855 Tiverton Dr., Los Angeles, CA, 90024, USA; Department of Surgery, The University of Chicago, 5841 S. Maryland Ave., Chicago, IL, 60637, USA

**Keywords:** finite element analysis, morphoelastic growth, imaging, simulation, meshing, geometry, shape, aortic dissection

## Abstract

The human aorta undergoes complex morphologic changes that indicate the evolution of disease. Finite element analysis enables the prediction of aortic pathologic states, but the absence of a biomechanical understanding hinders the applicability of this computational tool. We incorporate geometric information from computed tomography angiography (CTA) imaging scans into finite element analysis (FEA) to predict a trajectory of future geometries for four aortic disease patients. Through defining a geometric correspondence between two patient scans separated in time, a patient-specific FEA model can recreate the deformation of the aorta between the two time points, showing pathologic growth drives morphologic heterogeneity. A shape-size geometric feature space plotting the variance of the shape index versus the inverse square root of aortic surface area (δ𝒮 vs. 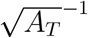) quantitatively demonstrates the simulated breakdown in aortic shape. An increase in δ𝒮 closely parallels the true geometric progression of aortic disease patients.

## 1. Introduction

Type B aortic dissection (TBAD) is a life-threatening vascular disease defined by the formation of a tear in the inner layer of the wall of the descending thoracic aorta, the largest artery in the human body. The incidence of TBAD is roughly 3 per 100, 000 people per year with a 5-year mortality rate of 30 − 40% [1]. The goal of clinical therapy is to slow disease progression and prevent complications, with treatments encompassing medication, endovascular repair, and surgical replacement [2]. Continued enlargement of the aorta, measured by an increase in cross-sectional-diameter, is a direct indicator of disease progression predictive of morbidity and mortality [3–5]. The yearly risk of aortic rupture is around 30% once the cross-sectional aortic diameter reaches 60 mm, and a key indication for surgical intervention is continued aortic dilitation, ocurring in over 70% of patients undergoing medical therapy and over 60% of patients with prior endovascular intervention [2, 6].

In addition to measuring cross-sectional expansion, the clinical literature has recognized that the aorta undergoes complex morphologic changes with disease progression [7]. Since size measurements are insufficient, current clinical guidelines aim to prevent morphologic changes indicative of aneurysmal degeneration and inferior outcomes [2, 8]. Multiple studies have identified aortic centerline tortuosity, mean centerline curvature, and cross-sectional eccentricity as risk factors for deleterious outcomes [9–14]. Others have applied statistical shape analysis (SSA), a mathematical approach to modeling shape variation in a population, to replicating clinical diagnoses, predicting surgical outcomes, and modeling rupture risk in aortic disease [15–18]. In a recent publication, we showed that projection of aortic anatomy into a space defined by size and the fluctuation in total curvature (*δK*), a measure of shape, differentiates aortas along the spectrum of growth and pathology [19].

Despite their prevalence in medical literature, statistical approaches are limited by the low prevalence of aortic diseases, coupled with the high risks of operative interventions, making the decision to treat a highly individualized one. Evidence from clinical trials relies on predicting future outcomes of an individual on the basis of outcomes in “similar” patients, yet finding a large-enough class of “similar” aortic disease patients is challenging [20]. This explains the paucity of high-quality clinical trials in the aortic disease literature. Furthermore, treatment algorithms must be interpretable to permit applicability for high-stakes medical decisions that rely upon pre-existing frameworks grounded in accepted practice. As such, lack of interpretability has been cited as a major obstacle to the adoption of novel artifical intelligence models in healthcare [21]. The absence of a clear mechanism driving such models severely hinders the medical community’s ability to personalize them [22].

Computational modeling, as an alternative predictive tool, may be better suited in addressing these challenges. Finite element analysis (FEA) and computational fluid dynamics (CFD) simulations have been applied to studying biomechanical mechanisms driving arterial diseases [23]. FEA models evolving aortic pathology as a mechanical process driven by physical forces. Past literature has centered on correlating predicted stresses with aneurysmal rupture risk, albeit with limited clincal application due to the inability of simulations to model underlying disease processes [23, 24]. Moreover, stress-based criteria are nearly impossible to translate because of the immense heterogeneity in local material properties of diseased aortas. While much progress has been made in elucidating non-linear and anisotropic aortic constitutive models, their homogenous application throughout aortic geometry remains a limitation. In our prior work, we showed using ideal shapes that varying local surface area expansion drives shape heterogeneity. We expand this methodology to real patient data and develop an imaging-informed finite element morphoelastic model of aortic shape evolution [19]. Similar work with morphoelasticity in aortic abdominal aneurysms modeled early simple uniform dilatation [25], and the stress analysis literature has failed to capture the heterogeneity that is the true hallmark of progressive aortic pathology.

We incorporate past geometric information into FEA to predict a trajectory of future geometries for aortic disease patients. Hypothesizing that pathologic growth drives morphologic heterogeneity, we aim to model the evolution of this breakdown. We show that non-uniform pathologic growth drives shape changes. Through defining a geometric correspondence between two patient geometries separated in time, a patient-specific FEA model can recreate the deformation of the aorta between the two time points.

## 2. Methods

### 2.1 Clinical Data Cohort

Four type B aortic dissection (TBAD) patients with pre-operative imaging are identified. Clinically, all four patients exhibit degenerating disease over several years that ultimately necessitated multiple operative interventions (see demographic information in Figure A.9). For this study, two preoperative scans are chosen to define the growth simulation for each patient. On average, scans were 2.6 *±* 1.2 years apart: 3.4 years for patient 1, 1.7 years for patient 2, 3.8 years for patient 3, and 1.5 years for patient 4.

Aortic geometries are obtained from computed tomography angiography (CTA) scans, as described in previous work [19]. Briefly, three-dimensional aortic models are extracted from imaging data with a workflow consisting of semi-automated segmentation, noise reduction, smoothing, isolation of the outer surface, and triangular surface meshing in the program Simpleware ScanIP (S-2021.06-SP1, Synopsys, Mountain View, CA) [19]. Triangular surface meshes for the 8 TBAD scans (2 per patient) consist of 38924*±*3278 elements.

The plots depicting real and simulated aortic shape vs. size for the four patients include a curve in the background representing empirically observed trends in physiologic and pathologic aortic shape evolution. This curve represents a power law (*δS* ∼ 𝓁^−3^, where 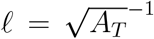) fit to a large patient cohort of 302 CTA scans, consisting of 131 scans of diseased aortas and 171 scans of non-pathologic aortas [19].

Geometries for FE simulations are obtained by discretizing the initial scan for the four TBAD patients as a solid linear tetrahedral mesh with a target of 3 elements through the thickness layer. Meshes consist of 307060 ± 47038 elements: 262593 for patient 1, 384166 for patient 2, 304793 for patient 3, and 276687 for patient 4.

All CTA images are obtained via the Human Imaging Research Office (HIRO) at the University of Chicago Medicine, an institutional imaging research core that provided HIPAA-compliant, deidentified DICOM data for patients requested for this study. A variety of scanners were used to collect radiographic information. All data collection and analysis is performed in accordance with the guidelines established by the Declaration of Helsinki and under institutional review board approval (IRB20-0653 and IRB21-0299).

### 2.2 Morphoelastic Growth Mapping

We develop a morphoelastic model linking geometry to the evolution of aortic dissection by incorporating patient-specific geometric changes into finite element analysis. We define a growth mapping ℳ that compares localized area changes between corresponding surface partitions on two patient geometries from different time points: *S* and *S*^*′*^. A data-derived growth field 𝒢 then defines a growth rate 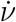 at each point on the aortic surface as the simulation input. Future local geometries are obtained from the simulation outputs.

Performing this growth mapping requires the definition of spatially corresponding surface partitions on two aortic geometries. First, rigid registration globally aligns the initial mesh *S* with the final mesh *S*^*′*^, followed by subdivision into *m* axial segments to allow for a stepwise non-rigid registration procedure that applies a non-rigid transformation 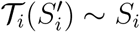 from each final segment 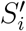 to initial segment *S*_*i*_. These collectively define a global transform 𝒯 ∼ (*S*^*′*^) *S* that spatially aligns the two geometries.

Once the geometries are spatially aligned, the initial geometry *S* is partitioned into *k* partitions, and the final geometry 𝒯 (*S*^*′*^) is equally partitioned.

Spatial correspondence defines a one-to-one mapping 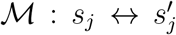 of the partitions, elements 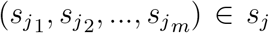 and 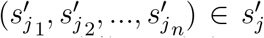, where *m ≠ n*. The area change of each partition 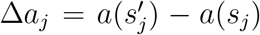 is used to calculate the per-partition growth rate, which is assigned to each element within the partition element set, each of which contains a set of Δ*a*_*j*_ to define the growth field 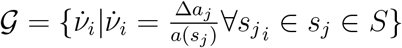. Finally, 𝒢 is smoothed to produce a continuous growth field that is projected onto the initial mesh geometry *S*. Figure 1 outlines the workflow for a hypothetical 2D example of two length-mismatched closed curves.

**Figure 1:**
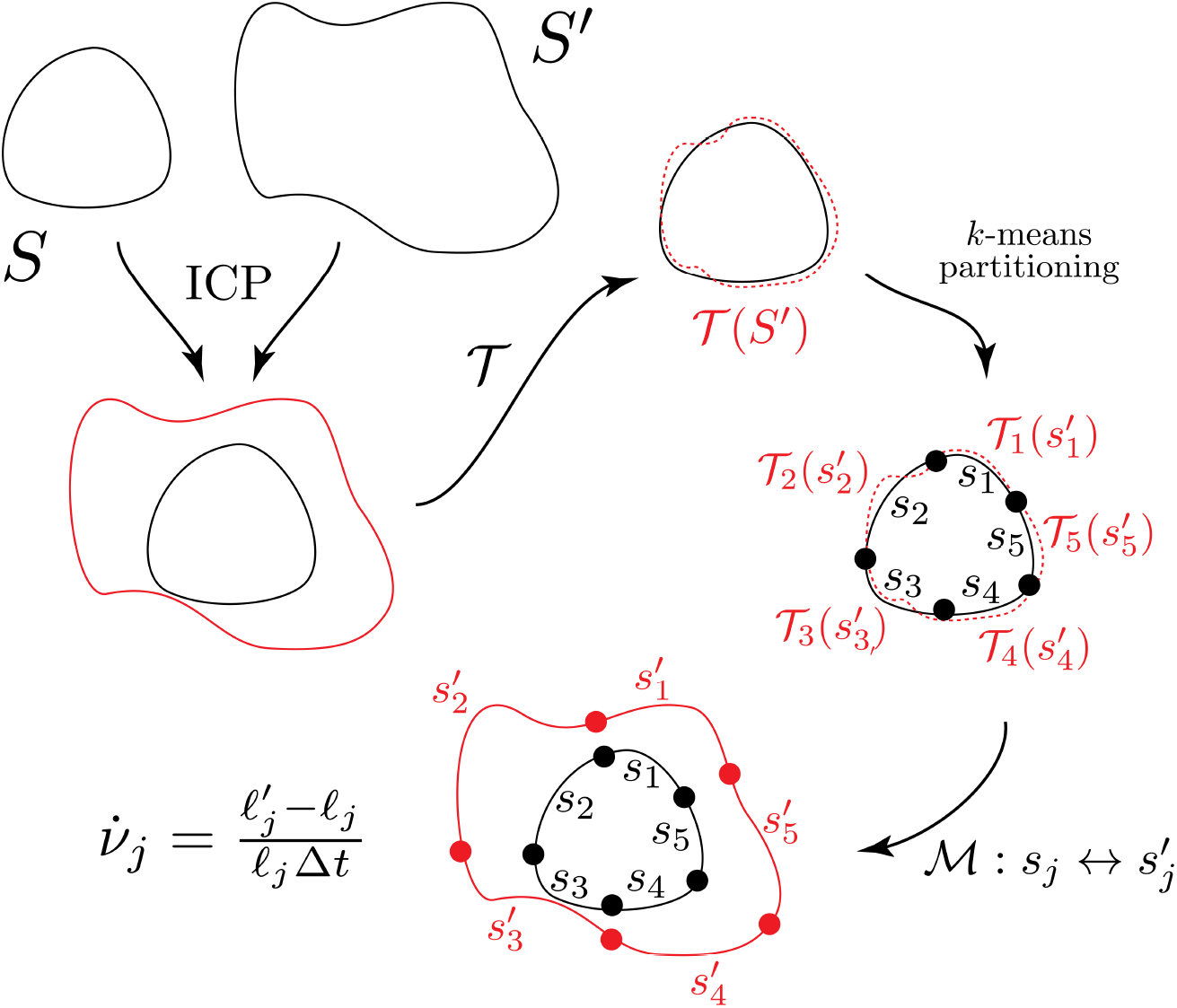
Schematic of our registration and local growth rate methodology in a simplified 2D case. The iterative point cloud (ICP) algorithm is used to spatially align *S* and *S*^*′*^, which are represented by two closed planar curves of different lengths and curvatures. The coherent point drift (CPD) algorithm defines a non-rigid transform 𝒯 such that 𝒯 (*S*^*′*^) is closely aligned with *S*. This spatial alignment allows for the *k*-means algorithm to define spatially corresponding partitions on the two curves such that each partition *s*_*i*_ on *S* is aligned with partition 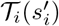 on *S*^*′*^. This allows for the definition of a one-to-one mapping 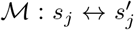 between partitions in the initial geometries. The partition growth rate 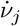 is calculated from the lengths 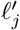 (on *S*^*′*^) and 𝓁_*j*_ (on *S*), respectively.

Anatomical correspondence between local regions of the two meshes is key to this method, which is why two registration procedures are necessary. This allows for calculation of local growth rates at the partition level via calculating the area change between corresponding partitions on the two geometries. Once growth rates are calculated at the partition level, smoothing ensures that they continuously vary over the aortic surface. This both minimizes discretization noise and permits the FE simulation to run successfully.

#### 2.2.1. Rigid Registration

The initial and final geometries are obtained from scans of a given patient at two different time points. The iterative closest point (ICP) algorithm, which minimizes the sum of squared differences, is used to physically align the meshes in coordinate space by translation and rotation of the initial geometry mesh onto the final geometry mesh, which remains fixed.

#### 2.2.2. Stepwise Alignment

Due to the extensive and nonuniform nature of aortic geometric evolution, rigid registration alone is insufficient to allow the calculation of local area changes. The misalignment hinders the calculation of a growth mapping, which requires the identification of small, corresponding regions between the two aortas whose difference in surface area reliably approximates the growth rate of the region. Non-rigid registration is used to model deformable motion by mapping correspondences between two sets of point clouds [26]. Given the many degrees of freedom arising from complex global shape changes, we constrain the problem by subdividing the each aortic surface along the centerline into *m* = 12 corresponding axial segments, running from the aortic sinus to the celiac trunk. The aorta is divided along its centerline to ensure that each segment represents a contiguous cross-section. For each mesh vertex *i*, a corresponding point on the centerline *k* is identified using the following algorithm:

1. Define a “search region” *R*_*i*_ of points on the centerline *C*(*t*) such that the arclength in each direction is 𝓁 = *L/m*, where *L* is the total arclength of the centerline.
2. Within *R*_*i*_, define a cone of selection *S*_*i*_ isolating nearly perpendicular centerline points as potential matches (set by a threshold parameter *ϕ* = 0.85). Let 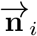 represent the vertex normal vector, constructed from the area-averaged normal vectors of surrounding mesh triangles. The vector from vertex *i* to a centerline point *j* is 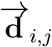.
3. Choose the point *k* in the cone of selection that is closest to vertex *i*.

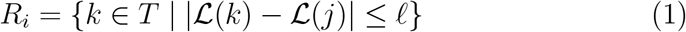

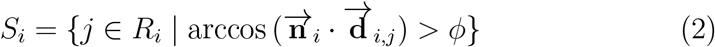

This subdivision simplifies the problem from aligning large, complex, and tortuous structures into aligning small cylindrical segments. We previously showed that aortas follow a universal linear scaling ℒ∼*c*𝓁 between centerline length ℒ and and cross-sectional size 𝓁, with scaling constant *c*∼*O*(10) according to multiple methods of measurement [19]. This motivates the selection of *m* = 12 segments.

Following subdivision, the vertices of each segment 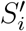 of the final scan are aligned with the vertices of the corresponding segment *S*_*i*_ on the first scan. The coherent point drift (CPD) algorithm models the alignment of two point sets by defining a non-rigid transformation 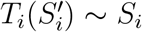 between a moving point set 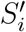 to a fixed set of data points *S*_*i*_ [26]. 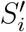 is modeled as Gaussian Mixture Model (GMM) centroids that are fit to the data points *S*_*i*_ via likelihood maximization. Regularization of the displacement field ensures that GMM centroids move coherently as a group, which preserves the topological structure of the point sets. This regularization helps preserve local geometric relations and ensure that mapped regions correspond in anatomical space [26].

We set a regularization parameter *λ* = 0.1, representing a trade-off between the goodness of maximum likelihood fit and regularization. The Expectation Maximization (EM) algorithm iteratively optimizes the GMM fit for a maximum of 50 iterations. A noise ratio of *w* = 0 indicates no expected noise in the point sets, and a parameter *β* = 4 sets the smoothness regularizer [26]. Figure 2 illustratess the alignment of three representative axial segments.

**Figure 2:**
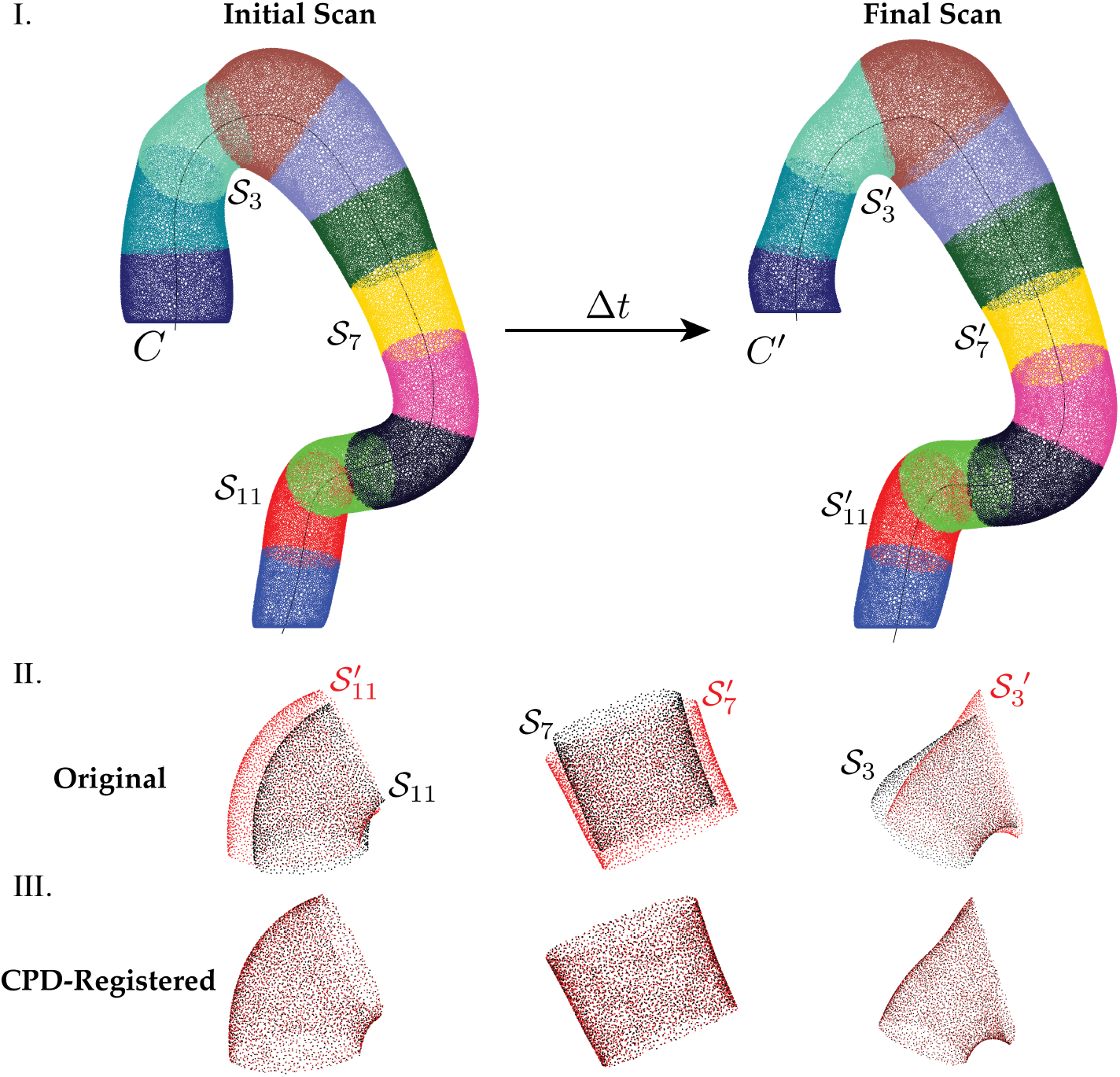
Complex aortic geometries are subdivided to allow for mapping local regions while preserving anatomic correspondence. Each axial segment has an approximately cylindrical shape that is well-suited for non-rigid registration using CPD. **I**. *(left)* mesh of the initial scan and *(right)* mesh of final scan used for geometric mapping. Each mesh is divided into 12 axial segments along the respective centerline. **II**. Original un-aligned meshes for three representative segments. The initial scan mesh is in black, and the final scan mesh is in red. **III**. Aligned meshes after 50 iterations of CPD.

#### 2.2.3 Mesh Partitioning

Following alignment, a one-to-one mapping 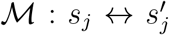 is defined between two sets of partitions *s*_*j*_ and 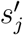 on the initial and final geometries, respectively. Mapping at the level of the partition permits for calculation of localized area changes between corresponding regions on the two geometries. This formulation both minimizes errors from mesh discretization and removes the requirment of mapping individual mesh elements, which would retrict this method to meshes of equal element count.

On the initial mesh *S*, vertices are grouped into *k* = 400 partitions using the *k*-means++ algorithm [27]. Matching partitions are obtained for the transformed second mesh 𝒯 (*S*^*′*^) with *k*-means++, here setting the initialization condition (i.e. starting centroids) as the cluster centroids from the first mesh. Centroid deformation is limited to a maximum of 5 iterations to allow for flexibility optimizing the partitioning of the second mesh while ensuring spatial centroid correspondence. Performing mesh partitioning on the transformation of the final scan 𝒯 (*S*^*′*^) ensures anatomical correspondance of partitions.

#### 2.2.4. Calculation of Per-Partition Growth Rates

A growth field 𝒢 is obtained from partition-level statistics. The growth rate of a partition is calculated as the proportional area change between corresponding regions with a linear time approximation:

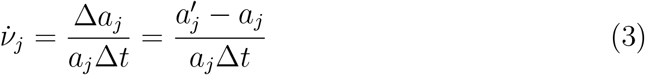

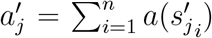 represents the area of a partition *j* in the second scan mesh, while 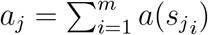 represents the area in the first mesh. Elements 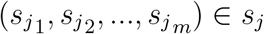 and elements 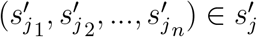, where *m* ≠ *n*.

Δ*t* represents the time difference between the two scans. Based on the hypothesis that positive area expansions drive morphologic changes in the aorta, constrained by the structure’s elastic continuity, we enforce nonnegative growth rates in the simulation by setting 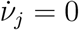 for regions with Δ*a*_*j*_ *<* 0. Note that area calculations are performed on the original surface meshes *S* and *S*^*′*^, and the transformed surface mesh 𝒯 (*S*^*′*^) is simply used to define the one-to-one map between ℳ corresponding partitions. The raw growth field is then

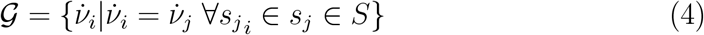

where 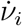 indicates the growth rate of an individual mesh element.

#### 2.2.5. Smoothing to a Continuous Growth Field

The partitioning procedure produces a discrete growth field with a single growth rate per partition. This is converted into a continuous field with a smoothly varying growth rate for finite element (FE) simulation. K-nearest neighbors (kNN) is used to find *n* nearest neighbors for each surface element and then set the growth rate of the element as the average growth rate of the nearest *n* neighbors, effectively transforming the growth rates into a smoothly varying field [28]. We set 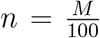, where *M* is the number of faces in the surface mesh.

The final step is to map the growth rates calculated on the surface triangulated mesh to the solid tetrahedral mesh that is used for simulation. Each tetrahedral element in the solid mesh is assigned the growth rate of the nearest triangular element in the corresponding surface mesh. Figure 3 illustrates the steps for transforming the partition-mapped growth rates into an FEA-ready continuous growth field.

**Figure 3:**
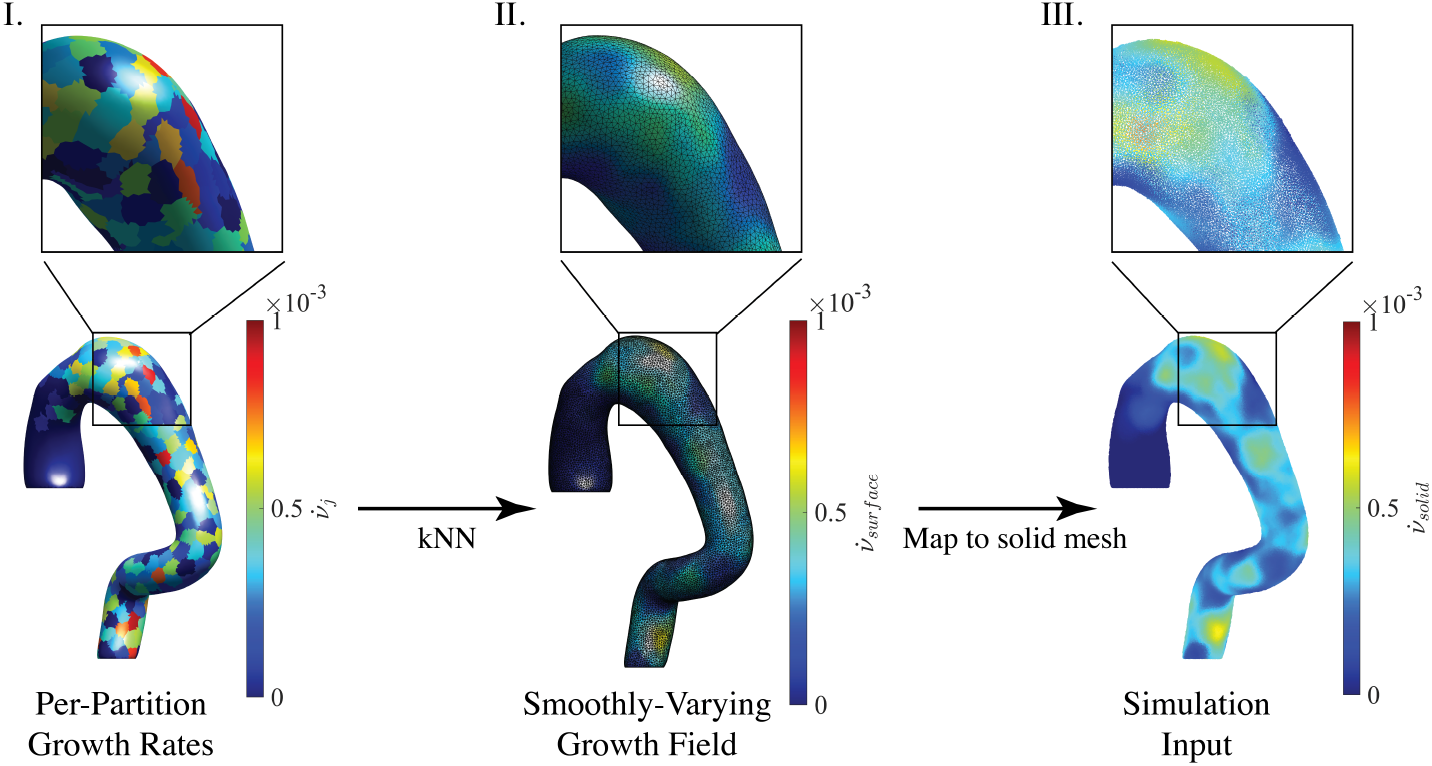
Generation of smoothly varying growth rates (in mm/day) for FEA simulation from aligned partitions. **I**. Growth rates are initially calculated for each aligned partition as 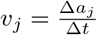 ; however, apparent noise (even with good alignment) necessitates smoothing to allow for successful simulation. **II**. Growth rates are smoothed using k-nearest neighbors (the triangular geometry of the surface mesh is shown). **III**. Surface growth rates are mapped to a solid mesh for FEA.

#### 2.2.6. Finite Element Simulation

Growth is computed using a morphoelastic model implemented in Abaqus (2021, Dassault Systèmes, Waltham, MA) that decomposes the deformation gradient *F* into an elastic contribution *F*_*e*_ and a growth contribution *F*_*g*_: *F* = *F*_*e*_*F*_*g*_ [29–31]. Adopting the assumption that *F*_*g*_ does not explicitly contribute to the free energy in previous work on computational modeling of multiplicative growth [29–31], a neo-Hookean (NH) strain energy function of the following form is used: *W* = *W* (*F, F*_*g*_) = *W*_*e*_(*F*_*e*_). When there is no growth, *F*_*g*_ = *I* and *F*_*e*_ = *F*, which represents a purely elastic deformation. With this decomposition and the NH strain energy, the stress can be derived and updated during the loading process according to the following relations:

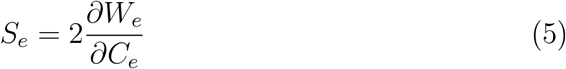

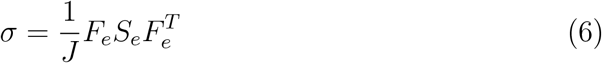

Where *S*_*e*_ and *σ* are the second Piola-Kirchhoff stress and Cauchy stress, respectively. Assuming isotropic growth with constant growth rate, the growth contribution *F*_*g*_ becomes:

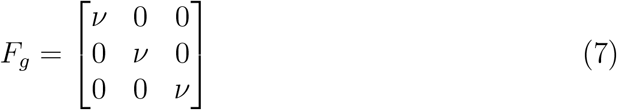

where the growth factor is 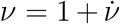, where 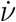 is the growth rate. To exclude the influence of material models, we universally employ a neo-hookean (NH) material with parameters *c*_10_ = *µ/*2 = 0.05 MPa and *D* = 2*/B* = 1 (*µ* is the shear modulus and *B* is the bulk modulus). The model is implemented with Abaqus Explicit solver and using the VUMAT subroutine [32, 33].

As previously stated, FE simulations employed solid linear tetrahedral meshes with a target of 3 elements through the thickness and 307060*±*47038 elements. The proximal and distal ends of the aorta, anatomically corresponding to the arotic sinus and the celiac trunk, are constrained using a fixed boundary condition (*u*_1_ = *u*_2_ = *u*_3_ = 0). The time length of the simulation corresponds to the time length between the scans, such that the total area change that was calculated earlier is recapitulated.

Fifty frames are selected from each simulation for geometric analysis. In order to analyze the deformed FE geometries, the 3D mesh must be transformed into a 2D surface. For each frame, elements corresponding to the outer surface are extracted from the solid mesh to form a triangular surface mesh with a consistent orientation of normal vectors.

### 2.3. Auto-Mesh Normalization

Our geometric analysis requires a constant mesh density between different geometries. The density of surface meshes directly derived from 3D imaging was optimized in our previous work at a target mesh element length of 0.5 mm [19]. This is significantly coarser than the densities of the FE meshes, which are refined to obtain simulation convergence. As stated in Section 2.1, surface meshes average at 38924 *±* 3278 elements, while solid meshes average at 307060 *±* 47038 elements.

Surface meshes extracted from FE simulations are compared to the constant target mesh density used in the geometric analysis of CT-derived data. This corresponds to an element length 𝓁_*t*_ = 0.5 mm. Assuming a triangular mesh consisting of equilateral triangles, the target element area is 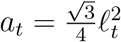. The target density is obtained via an automated mesh density normalization proceedure consisting of decimation and smoothing. Element count is reduced by progressively merging mesh elements until reaching the target element area. This is defined with a target ratio of preserved faces *r* = *a*_*m*_*/a*_*t*_, where *a*_*m*_ is the mean element area of the initial mesh. Then, a smoothing filter controlled by a strength parameter *f* = 0.5 is applied for 5 iterations to optimize the surface topology following the decimation procedure. This is followed by noise reduction using a volume-preserving Laplacian smoothing filter with parameters *λ* = 0.5 and *λ*_border_ = 0.5, normalized by element size, applied for 10 iterations. This mesh normalization procedure is performed in Blender (2018, Blender Foundation, Amsterdam, NL).

### 2.4. Geometric Analysis

Simulated geometries are compared to real CT image-derived geometries via quantification of shape and size metrics. The Rusinkiewicz algorithm calculates the per-vertex shape operator 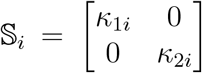 as the weighted average of the shape operators of immediately adjacent faces [19]. Mesh geometry is quantified from the per-vertex principal curvatures *κ*_1_ and *κ*_2_. Size is parameterized as the total surface area *A*_*T*_ . While the fluctuation in total curvature (*δK*) was used in previous work, here we use the Koenderink shape index 𝒮:

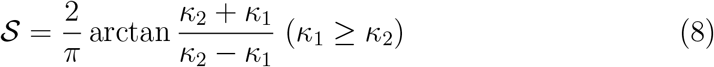

𝒮 is a scale-invariant shape operator which parameterizes the per-vertex principal curvatures onto a 1-dimensional disc in (*κ*_1_, *κ*_2_)-space [34, 35]. The bounded nature of the function (𝒮 ∈ [−1, +1]) maps all geometries, except for planar surfaces, onto an intuitive scale ranging from a concave spherical surface (𝒮 = −1) to a convex spherical surface (𝒮 = +1) [34].

The surface variance in shape index *δ 𝒮* captures the aorta’s deformity in shape. *δ 𝒮* is calculated for a geometry as the second moment (*µ*_2_) of a probability mass function, constructed as a histogram of the vertex-level shape index *s*_*i*_, weighted by the corresponding vertex voronoi area *a*_*i*_ (a weighted average of the areas of neighboring faces) [34, 36]:

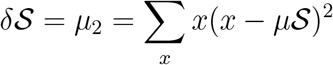

All calculation is performed in Matlab (2023b, Mathworks, Natick, MA). Aortic geometries are extracted from patient CTA imaging scans using a custom workflow in Simpleware ScanIP (S-2021.06-SP1, Synopsys, Mountain View, CA) including semi-automated segmentation, smoothing, outer surface isolation, and triangular surface meshing. Smoothing consists of an optimized five-step procedure includion geometric dilation, a mean filter, a median filter, a recursive Gaussian blur filter, and geometric erosion. Noise reduction is performed by characterizing a single aorta as 15 triangular meshes of varying density and smoothing intensity, then averaging the resulting 𝒮 *δ* values [19]. Details of the preceeding have been previously described [19].

## 3. Results

The growth-based FE model is tested on four examples of evolving TBAD. A growth field 𝒢 is constructed for each patient by mapping changes in the area of corresponding surface partitions between two scans, collected over an average timespan of 2.6 years (Figure 4). As shown in Figure 5, 𝒢 is smoothed to a continuous growth field to serve as the input for the FEA simulation. The simulation models the geometric evolution from the first scan according to the local growth field (Figure 6), with the goal of capturing the geometry at the second time point. Figure 7 compares the first and second imaging scans with the initial and final geometries from the simulation. Visual inspection allows for comparison of the actual imaging-derived final scan with the predicted simulation-derived final scan, showing a high degree of similarity.

**Figure 4:**
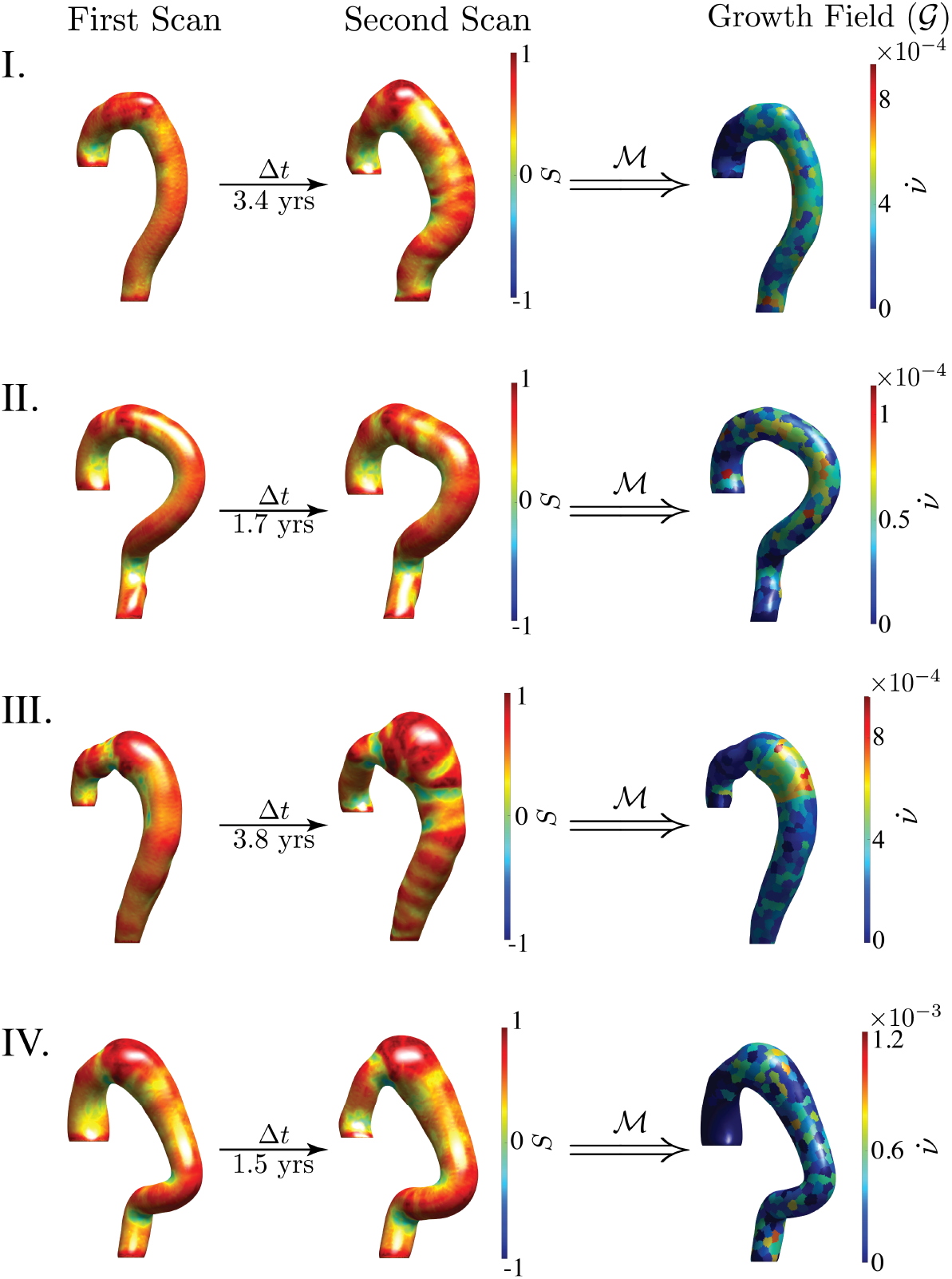
Heatmaps outlining growth mapping capturing geometric change. The first and second imaging scans represent two aortic geometries. As described in Section 2.2, our morphoelastic model defines a growth mapping ℳ that compares local area changes between corresponding surface partitions on the two patient geometries. This produces a data-derived growth field 𝒢 that defines a growth rate 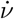 (in mm/day) on each point of the aortic surface, Δ*t* = 3.4 years (I), 1.7 years (II), 3.8 years (III), 1.5 years (IV).

**Figure 5:**
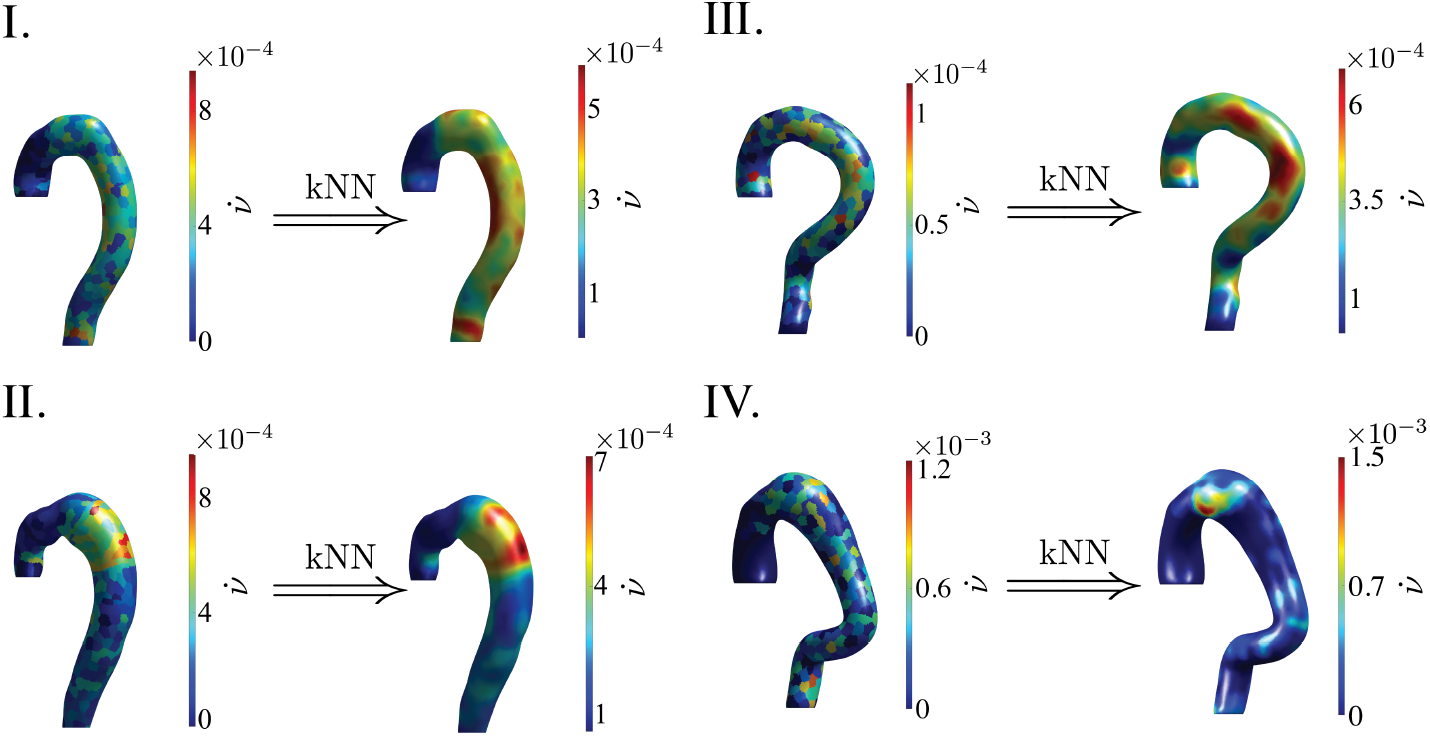
Smoothing to produce a continuous growth field. The growth field 𝒢 must be smoothed using a k-nearest-neighbors (kNN) algorithm to obtain a smoothly varying growth rate (in mm/day) that is suitable for FE simulation.

**Figure 6:**
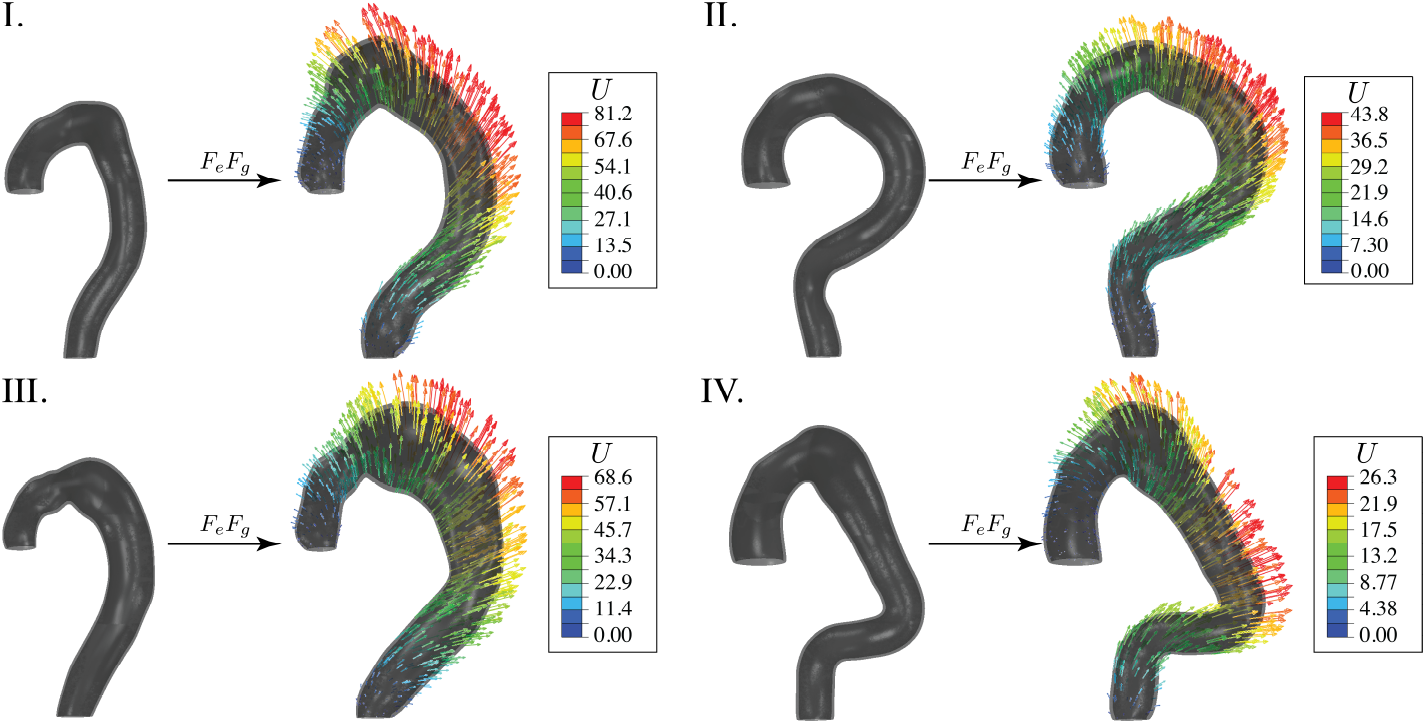
FE simulation deformation. The FE simulation incorporates a morphoelastic model that decomposes the deformation gradient *F* into an elastic component *F*_*e*_ and growth component *F*_*g*_, where *F*_*g*_ is defined by the smoothed growth field 𝒢. The deformation vectors *U* indicate the geometric change from the initial scan in millimeters (mm).

**Figure 7:**
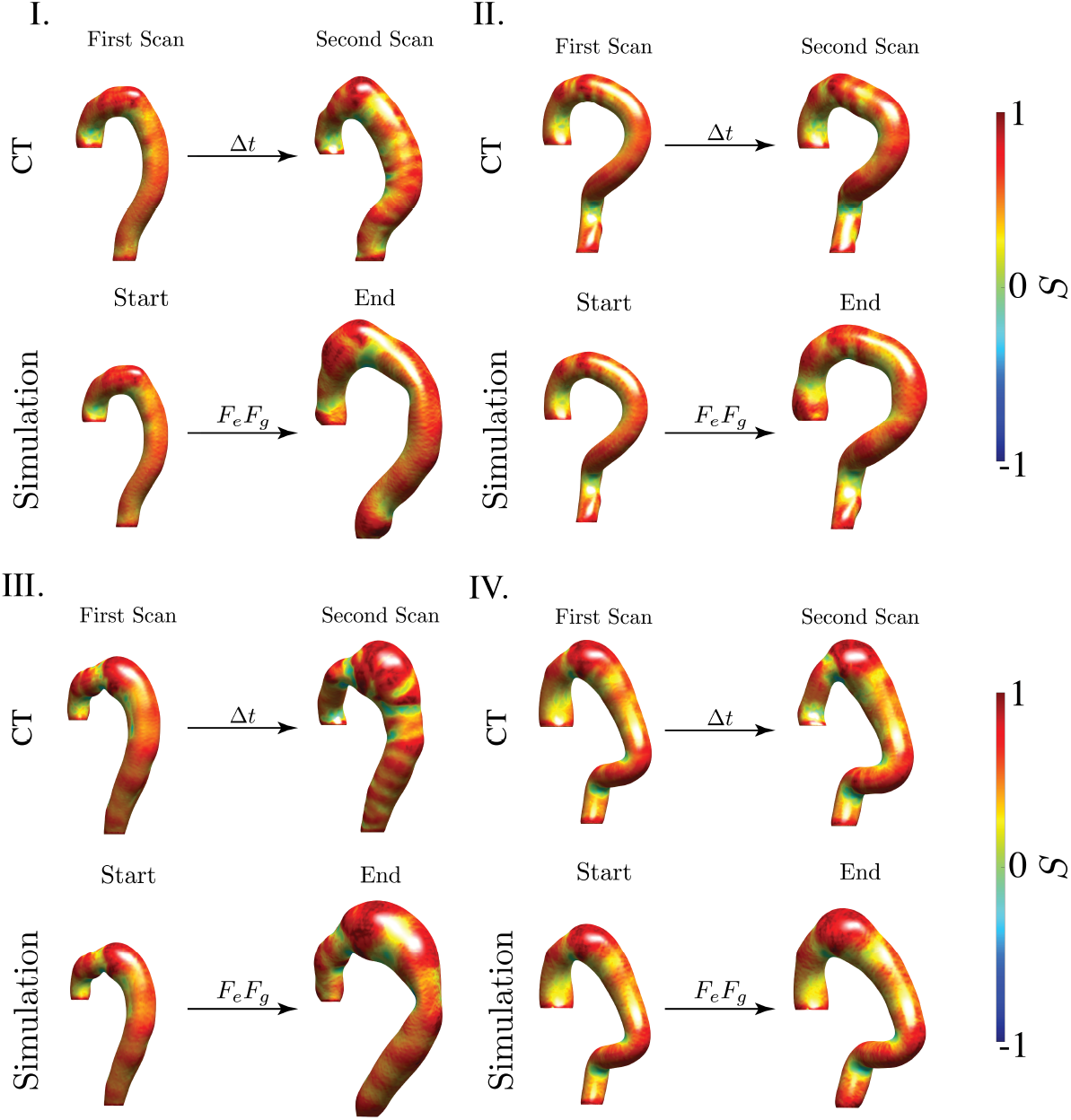
Comparison of patient scans with simulation-derived aortic geometries. The FE simulation predicts aortic deformation from the patient’s first scan and the mapped growth field 𝒢. For each patient, the top row shows the CT-derived geometries, while the bottom row indicates the FE simulation-derived geometries. Note that all simulation-derived surface meshes are normalized to a constant mesh density, as described in section 2.3.

Figure 8 provides quantitative comparisons of the trajectories of each patient’s aorta, as captured in the imaging scans, with the corresponding computational FEA simulation. In each figure, a dotted curve represents a power law fit 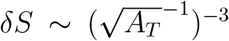 to a large patient cohort depicting the distribution of aortic geometries across the spectrum of growth and pathology [19]. Shapes connected by the black lines indicate all of the imaging scans for each patient of interest, organized over time. The large blue and red squares specifically indicate the initial and final scans used to generate the growth mapping for the FEA simulation, respectively, while the small black circles show additional scans. Hollow circles with the green-to-yellow color gradient show the geometric trajectory through the course of the FEA simulation. Appendix A.2 plots comparative results with a uniform growth field.

**Figure 8:**
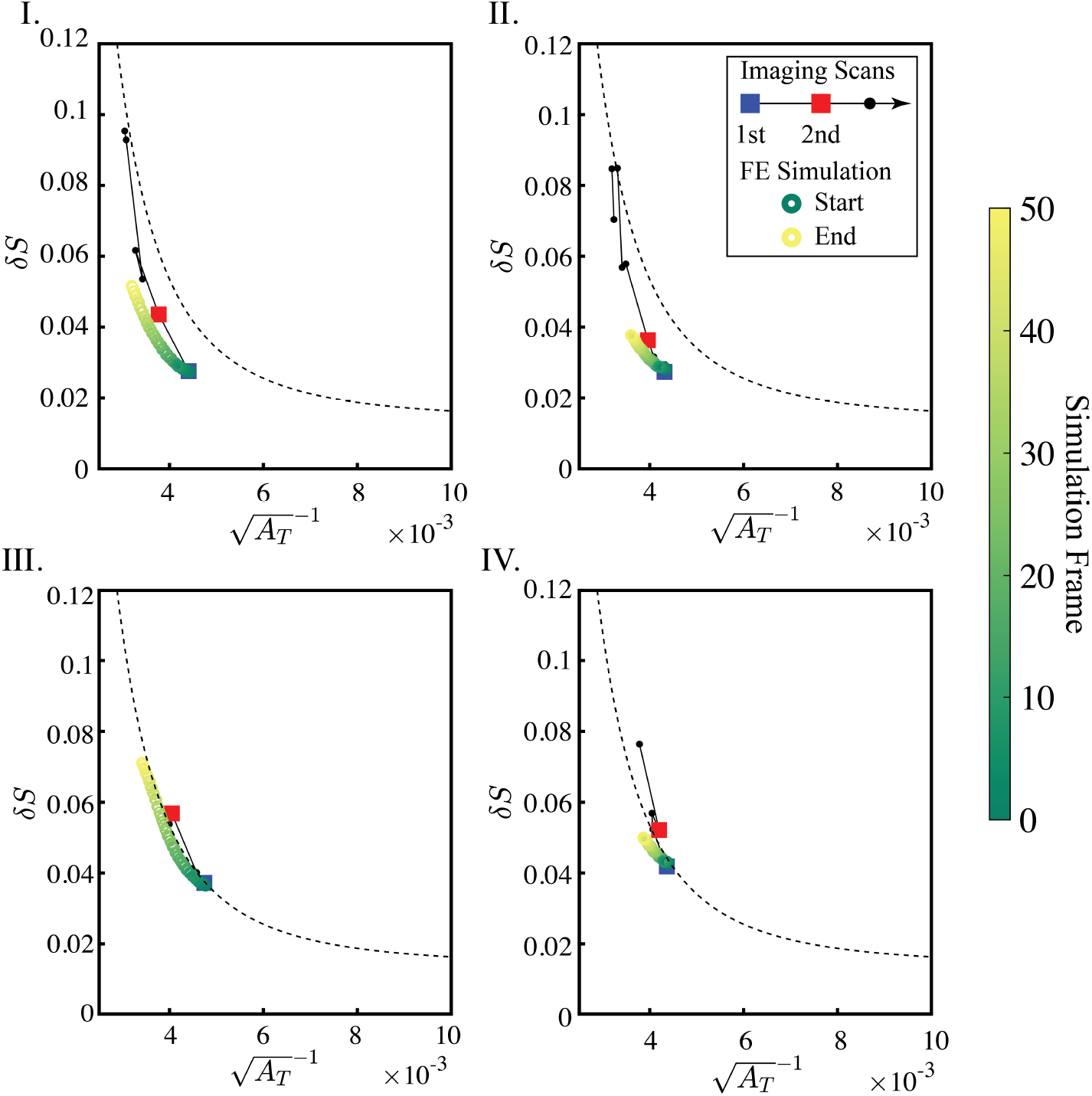
Comparison of true geometric evolution and FEA predicted geometric evolution in the 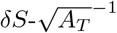 feature space. The green-to-yellow gradient indicates geometry at each simulation frame. The shapes connected with a black line show real patient geometries at different time points. Large blue and red squares indicate the initial and final patient scans used to define the geometric mapping for the FEA simulation, respectively. Small black circles indicate the patient’s remaining imaging scans. The dotted curve represents geometric trends across a range of aortas from a large patient cohort [19].

## 4 Discussion

Patients suffering from aortic disease, dissections or aneurysms, are often clinically followed with serial CT imaging. We recently developed a scale-invariant measure of aortic shape that fully characterizes, in a reduced twodimensional feature space, aortic morphology along the normal-pathologic axis, when combined with aortic size [19]. This prior work was purely geometric and lacked any connection to the underlying biomechanical mechanisms that drive aortic morphology. In this paper, we show that the source of the morphologic shape signal (*δS* and *δK*), whose hallmark is the loss of aortic shape self-similarity over time and relative to the general population, is embedded in the spatial gradients of local aortic growth (𝒢). We implemented a global and local co-registration methodology between two patient-specific aortic surface geometries obtained from CT scans at different time points (Δ*t*). The underlying hypothesis that a pathologic growth process drives morphologic heterogeneity of the aorta is demonstrated by the high degree of similarity between real and simulation-predicted aortic geometric trajectories, both in visual 3D space (Figure 7) and the *δ 𝒮* vs. 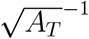 feature space. Interestingly, all four simulations exhibit a trend of increasing aortic surface area (decreasing 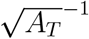) with increasing shape deformity (increasing *δ 𝒮*). This clearly matches a previously described aortic shape scaling law coupling increasing size with increasing shape deformity [19].

Our simulations are carried out using an explicit solver, as such no global energy minimization is imposed on the time evolution of the solutions. Growth enters the elasticity problem locally per element (*F*_*gj*_) by modifying the elastic deformation gradient 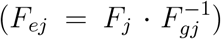 which updates the local stress 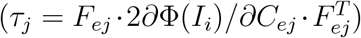. The updated stress *τ*_*j*_ is used to solve the compatibility problem assuring balance of forces and momenta globally [37, 38]. This global stability problem is sensitive to overall aortic shape.

A remarkable result is the overall smooth and stable deformation field obtained in our FEA simulations (see Figure 6), given the initial non-linear mixed-shell geometry and the non-linear loading through growth. Mixed shells are well-known to undergo complex elastic instabilities relative to homogenous plates and shells [39, 40]. By definition, mixed shells have spatial gradients in Gaussian curvature, *κ*_*g*_. Internal geometric boundary layers occur along crown lines where *κ*_*g*_ = 0 and are control regions for mixed shell stability, often leading lead to unpredictable and sudden buckling/wrinkling [39, 40].

The aortic geometries we study are all highly non-linear mixed shells with heterogeneously distributed hyperbolic, spherical, and parabolic regions. Figure 4 shows spatial gradients of 𝒮 ∼ *K* ∼ *κ*_*G*_ *×* area (see Appendix A.3 for details), with 𝒮 ∼ 0 corresponding to parabolic regions, 𝒮 ∼ *±*0.5 to hyperbolic regions, and 𝒮 ∼ *±*1 to spherical regions. Crown lines (𝒮∼0, green) are seen to course through all four initial scan geometries and become more pronounced in the aortas which deform extensively in the second scan (see Figure 4 I and III.). The correspondence between the CT and simulation geometries is poorest around the crown lines (compare second scan and simulation end panels in Figure 7). While the simulations fail to capture the appearance of these internal geometric boundary layers, it is remarkable that they correctly capture the evolution of the nearby spherical regions while maintaining global shell stability.

The overall stability of the deformed solutions is further remarkable given the known propensity for heterogenous growth to trigger elastic instabilities in simpler geometrically homogenous shells (spheres and cylinders) [37, 38]. These instabilities are known to occur due to the development of growth induced residual stress. The later develops in situations where a compatible zero-stress solution does not exists for the given geometry after growth despite the lack of external boundary conditions or loads.

Detailed studies of growing mixed shells are lacking [37, 38]. Our results, however, show that the particular growth fields that develop during the aneurysmal evolution of an aortic dissection are overall highly robust from the standpoint of shell elastic stability. This somewhat surprising finding leads us to hypothesize that the spatially non-homgenous growth field 𝒢 encodes a unique mechanically stable solution which minimizes residual stresses for non-linear mixed-shell geometries.

As a numerical solver for differential equations, FEA can computationally predict aortic shapes at any given future point, extrapolating beyond the initial scans used to define the growth field. Once the growth contribution *F*_*g*_ to the total deformation gradient *F* is defined, continued growth may be modeled assuming that *F*_*g*_ is constant. As such, a potential clinical application is that one may produce multiple predictions of shape at future time points and compare each predicted shape to the *δS* decision boundary. This would allow one to predict if a surgical operation may be successful at any arbitrary time point. Geometric characterization on predicted future geometries is an especially important application given the lack of consensus in the clnical literature on timing of CTA imaging for TBAD, as well as the limitations in obtaining frequent imaging due to poor patient compliance and socioeconomic barriers [41, 42].

While the assumption of a constant *F*_*g*_ may not be fully acurrate given the evolution of a disease process of long time scales, it paints a fundamental distinction between modeling approaches (like FEA) and the direct predictive approaches common in the statistical literature. While a predictive statistical model, such as a supervised learning method, would only be trained to predict shapes within the given time frame, FEA naturally allows for predictions of future geometries via extrapolation of the simulation. This coupled approach also carries the benefit of explainability, by which FEA generates concete structure predictions that are projected onto an interpretable geometric feature space (*δ𝒮* as a shape measure vs. 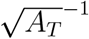 as a size measure). This is a major benefit over concurrent data-driven approaches in the literature, including statistical shape analysis (which derives from principal component analysis), supervised learning methods (including support vector machines and and decision trees), and convolutional neural networks [16, 43, 44].

### 4.1 Limitations

A major limitation of this work is that the growth field, while spatially varying to match the deformation between two given scans, is temporally constant (derived via linear approximation). We chose this formulation because of our interest in how linearly applied spatial gradients of growth may lead to nonlinear geometric evolution. Nonetheless, it is well known that biological structures are inherently dynamic, especially in pathologic conditions [29]. As such, there is an expectation that simulation accuracy will decrease over longer time scales, which makes extrapolation more uncertain. A time-dependent growth field may be defined by calculating the growth component of the deformation gradient *F*_*g*_ using a nonlinear approximation from multiple CTA scans. This may allow for the definition of a function *F*_*g*_(*t*) describing how the growth rate at a particular point changes over time. If such a function has a simple analytical form, then *F*_*g*_(*t*) can be extrapolated in time, potentially better predicting future aortic shape deformation.

A second limitation is that our method of geometric analysis, 𝒮*δ*, is a global shape descriptor that characterizes the variance of the shape index distribution across the entire aortic surface, thus ommiting local information. Changes in 𝒮 are treated as equal regardless of where they occur on the aortic surface, missing clinical knowledge that geometric degeneration in certain regions are highly correlative of certain aortic pathologies and complications [45, 46].

Finally, this was a retrospective study conducted at a single medical center on a pilot cohort of four patients. Future work should first expand to a larger cohort size with a wider degree of aortic pathologies, clinical course, and time between simulation input scans. This may reveal potential drawbacks or necessary modifications to our method in different application settings, as well as clarify the relationship between time extrapolation and prediction error. Prospective studies will provide higher-level clinical evidence, whereby a patient’s future geometric shapes at different time points may be predicted from two initial visits and then validated with follow-up imaging. Furthermore, external validation at multiple care settings is crucial before implementing any predictive model in clinical practice [47].

## 5. Conclusion

Finite element analysis (FEA) driven by curvature-mapped morphoelastic growth shows promising trends in modeling aortic shape evolution of an individual patient’s anatomy, demonstrating an intrinsic linkage between aortic geometry and mechanical stability. Given that past mechanical changes in the aorta provide insight into future evolution under a constant disease process, our curvature-mapped growth model may be further refined for application to predicting future geometric changes. Individualized prediction of a patient’s future aortic geometry carries the potential to improve aortic disease management by better informing timing of surveillance imaging and surgical interventions. In this paper, we show that the source of the morphologic shape signal (*δS*), which has been shown to be the hallmark of aortic pathology [19], is embedded in the spatial gradients of local aortic growth (𝒢). Furthermore, our results support the hypothesis that the spatially non-homgenous growth field 𝒢 encodes a unique mechanically stable solution which minimizes residual stresses for non-linear mixed-shell geometries.

## Acknowledgments

We acknowledge the support of the National Institutes of Health, NHLBI, R01HL159205 to LP. The Center for Research Informatics is funded by the Biological Science Division at The University of Chicago with additional funding provided by the Institute for Translational Medicine, CTSA grant number ULITR000430 from NIH. The funders had no role in study design, data collection and analysis, decision to publish, or preparation of the manuscript.

## Appendix A. Appendix

### Appendix A.1. Demographic Information

**Figure A.9:**
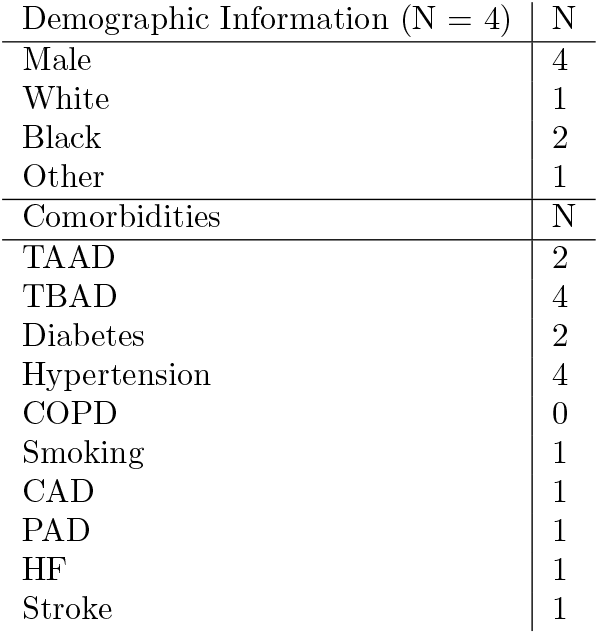
Patient Information. TAAD: Type A aortic dissection, TBAD: Type B aortic dissection, COPD: Chronic obstructive pulmonary disease, CAD: Coronary artery disease, PAD: peripheral artery disease, HF: heart failure.

#### Appendix A.2. Simulation of a Uniform Growth Field

The results presented in Figure 8, based on the geometry-mapped growth field *G*, are compared to an alternative uniformely-mapped growth field 𝒢_0_:

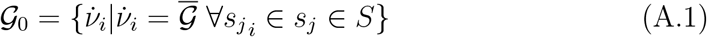

This sets a uniform isotropic growth rate 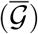 that is the mean of the mapped element growth rates. Figure A.10, which plots the results of this set of simulations, demonstrates that the shape trajectory achieved by uniform growth does not mirror patients’ true aortic shape changes. This shows that the co-registered and mapped gradients of local aortic growth (𝒢) were necessary to produce the trend of increasing aortic surface area with increasing shape deformity that matches observed aortic shape evolution [19].

**Figure A.10:**
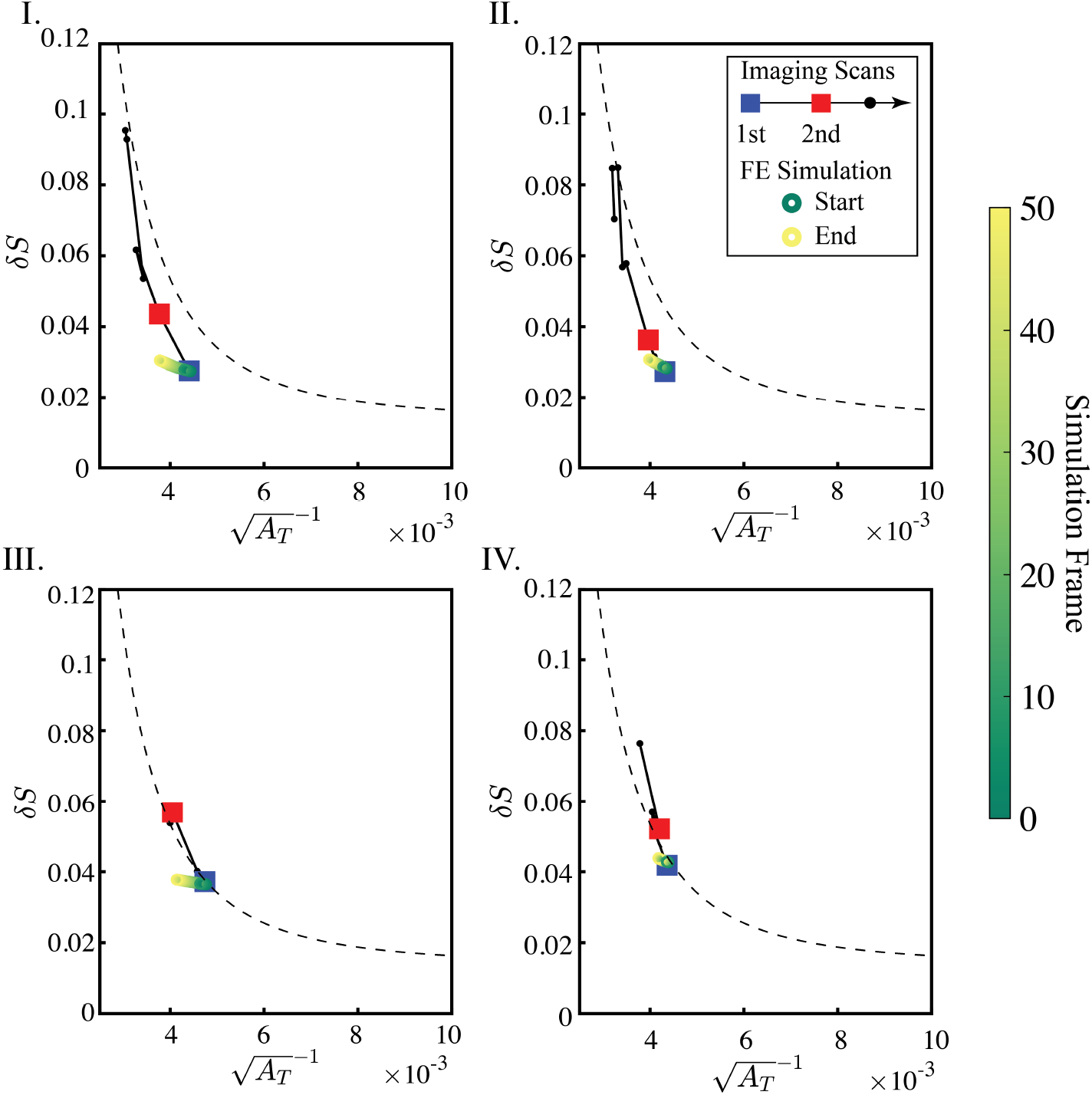
Comparison of true geometric evolution and FEA predicted geometric evolution for a uniform growth model in the 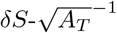 feature space. The green-to-yellow gradient indicates geometry at each simulation frame. The shapes connected with a black line show real patient geometries at different time points. Large blue and red squares indicate the initial and final patient scans used to define the geometric mapping for the FEA simulation, respectively. Small black circles indicate the patient’s remaining imaging scans. The dotted curve represents geometric trends across a range of aortas from a large patient cohort [19]. With uniform growth, unlike with the mapped morphoelastic model demonstrated in Figure 8, simulation trajectories do not align with the imaging scans.

#### Appendix A.3. Mathematical Connection Between Total Curvature and the Shape Index

We previously showed that divergence in the fluctuation in total curvature *δK*, which characterizes heterogeneous morphologic evolution of aortic surfaces through local shape changes, is highly predictive of surgical failures, such that a linear decision boundary in the *δK* vs.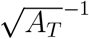 space may delinate successful vs. unsuccessful operations with high accuracy [19].

Total curvature (*K*) is obtained through mapping normal vectors on the external aortic surface onto the unit sphere **S**^2^, operationally implemented through integration of the per-vertex Gaussian curvature over a region of approximately constant curvature: 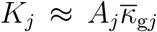 (*A*_*j*_ is the surface area of the region and 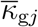 is the mean Gaussian curvature in the region) [19]. Due the operational discretization of the aortic surface into a triangular surface mesh, as well as the unbounded nature of *κ*_g_, the calculation of *K* is subject to numerical errors.

Unlike total curvature, the shape index is an empirical function with intuitive parameterization of surface shapes [35]. It is bounded in range (𝒮 ∈ [−1, 1]), with concave shapes populating 𝒮 ∈ [−1, 0) and convex shapes populating 𝒮 ∈ (0, 1]. The extreme ends represent a sphere (𝒮= −1 and 𝒮 = +1), while cylindrical shapes are at 𝒮= *±*0.5. These properties not only make well-defined and computationally simple to calculate, but also resistant to numerical errors [34].

*K* derives from extensive mathematical theory and is generally applicable beyond explicit mesh-based calculations for two-dimensional surfaces in R^3^. However, the increased noise sensitivity of *K* complicates calculations with geometries resulting from numerical simulations. Since both functions characterize shape in a size-independent matter and are mathematically connected, we decided that 𝒮 would be more appropriate for this study.

Drawing on *δK*’s characterization of post-operative surgical success for TBAD, it is possible define of a corresponding decision boundary in the *δ*𝒮 vs.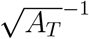 space representing a critical point where endovascular intervention may no longer be beneficial. Integrating the capabilities geometric method with the generative nature of numerical simulations allows us to postulate synergistic benefits in improving clinical predictions for aortic pathologies..

One can postulate a mathematical connection between *K* and 𝒮, which both calculate shape in a size-invariant manner:

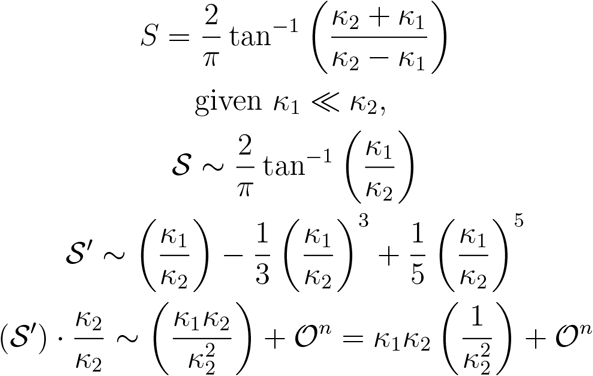

The first order term is dimensionally related to the total curvature *K* ∼ *κ*_*G*_ *×* area; the area over which the Gaussian curvature is integrated (𝓁^2^) is set by the smallest local dimension: 𝓁∼1*/κ*_2_. The power expansion tells us that *K* contains the same information as *S* when one local dimension is much smaller than the other (*R*_2_ ≪ *R*_1_), as in cylindrical geometries: 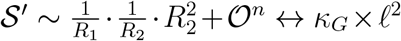, then the following must hold 𝒪 (*R*_2_) ∼ *𝒪* (𝓁^2^)

To first order, the shape index is a very specific way of integrating the local Gaussian curvature over an area set by the inverse of the largest local principle curvature. This new insight into the shape index connects it to the theoretical framework of differential geometry and provides mathematical backing to complement its practical utility as a bounded and intuitive shape function.

